# Differential surface competition and biofilm invasion strategies of *Pseudomonas aeruginosa* PA14 and PA01

**DOI:** 10.1101/2021.05.17.444588

**Authors:** Stefan Katharios-Lanwermeyer, Swetha Kassety, Carey D. Nadell, George A. O’Toole

**Author notes:** To whom correspondence should be addressed Carey D. Nadell, Rm 326 Class of 1976 Life Science Center, 78 College St., Dartmouth, Department of Biological Sciences, Hanover, NH 03755, George A O’Toole, Rm 202 Remsen Building, 66 College Street, Geisel School of Medicine at Dartmouth, Hanover, NH 03755. These authors contributed equally to this work (Stefan Katharios-Lanwermeyer, Swetha Kassety).

## Abstract

*Pseudomonas aeruginosa* strains PA14 and PAO1 are among the two best characterized model organisms used to study the mechanisms of biofilm formation, while also representing two distinct lineages of *P. aeruginosa*. Our previous work showed that *P. aeruginosa* PA14 and PAO1 use distinct strategies to initiate biofilm growth. Using differentially-labeled strains and microfluidic devices, we show that PAO1 can outcompete PA14 in a head-to-head competition during early colonization of a surface, can do so in constant and perturbed environments, that this advantage is specific to biofilm growth and requires production of the Psl polysaccharide. In contrast, the *P. aeruginosa* PA14 exhibits a competitive fitness advantage when invading a pre-formed biofilm and is better able to tolerate starvation than PAO1 in the biofilm context. These data support the model that while *P. aeruginosa* PAO1 and PA14 are both able to effectively colonize surfaces, these strains use distinct strategies that are advantageous under different environmental settings.

**Importance:** Recent studies indicate that *P. aeruginosa* PAO1 and PA14 use distinct strategies to initiate biofilm formation, with PAO1 committing to the surface through a processive mode of attachment, while PA14 uses a non-processive surface engagement strategy. We investigated whether their respective colonization strategies impact their ability to effectively compete under different biofilm-forming regimes. Our work shows that these different strategies do indeed impact how these strains colonize the surface: PAO1 dominates during colonization of a naïve surface, while PA14 is more effective in colonizing a pre-formed biofilm or withstanding starvation conditions. These data suggest that even for very similar microbes there may be distinct strategies to successfully colonize and persist on surfaces during the biofilm life cycle.

## Introduction

Biofilms are surface-attached microbial communities that mediate long-term growth and persistence on substrates as varied as vegetable produce and indwelling medical devices (1, 2). Biofilms are typically polymicrobial in nature and exhibit an increased tolerance to antimicrobial agents, predation and reactive oxygen species (3–5). The transition from planktonic to biofilm modes of growth is complex and uses signaling systems initiated by surface engagement that have downstream consequences for biofilm formation (6–8). Our understanding of biofilm formation comes in part from the study of *Pseudomonas aeruginosa*, a Gram-negative bacterium that forms prolific biofilms in aquatic environments, soils and clinical settings (9–12). To form a biofilm, motile bacteria such as *P. aeruginosa* use appendages including flagella and pili to mediate early surface engagement (13, 14). Surface colonization and growth during biofilm formation are also linked to extracellular polysaccharide (EPS) production, which provides cell-to-surface and cell-cell adhesion (15). EPS is required for early attachment events, particularly in the PAO1 lineage, as well as microcolony development; this matrix material also plays a key role in providing the structural integrity required for mature biofilms to form and persist (16, 17).

Current evidence suggests that *P. aeruginosa* use two distinct signaling systems to initiate formation of a biofilm (18, 19): the second messenger c-di-GMP, which is regulated through the Wsp system (20–22), and the second messenger c-AMP (cAMP), which is regulated through the Pil-Chp/Vfr system (23, 24). For *P. aeruginosa* PAO1, upon surface attachment, membrane-bound WspA is thought to behave as a sensor of surface contact, contributing to the phosphorylation and activation of the diguanylate cyclase WspR (22, 25) and resulting in increased c-di-GMP synthesis (26). In contrast, for *P. aeruginosa* PA14, signaling via the Pil-Chp system is initiated through type IV pili (TFP) engaging a surface, which in turn transmits this surface signal to the adenylate cyclase, CyaB. The activation of CyaB increases concentrations of cAMP, resulting in production and secretion of cell surface-localized PilY1. The PilY1 protein participates in the activation of diguanylate cyclase SadC by an unknown mechanism to enhance c-di-GMP production (24, 27–29). Thus, both of these pathways ultimately increase c-di-GMP level and enhance biofilm formation.

The two common laboratory strains of *P. aeruginosa*, abbreviated PAO1 and PA14, represent two distinct lineages based on whole genome phylogenetic analysis (30). PAO1 and PA14 preferentially use the WspR and Pil-Chip signaling pathways to enhance EPS production and repress surface motility, respectively, during early events in biofilm formation (24, 31). Previously, we have suggested that PAO1 uses the Wsp system to facilitate a processive regime of attachment that results in an early commitment to a surface (19). In contrast, PA14 is non-processive, such that cells are less committed after initial surface engagement while their progeny are primed for subsequent reattachment through a cAMP-dependent signaling pathway (19, 32).

The respective strategies by which PAO1 and PA14 are able to attach to a surface provide a useful model system by which we can test how initial colonization strategies influence competitiveness in the biofilm context when both strains are grown together, and thus compete for space and nutrients. The different surface-attachment strategies of PAO1 and PA14 may also provide insights in understanding how various bacterial genera, species and strains cooperate and/or compete to colonize surfaces in polymicrobial environments.

## Results

### PAO1 outcompetes PA14 in dual-strain biofilms

To test if the colonization strategies used by *P. aeruginosa* PA14 and *P. aeruginosa* PAO1 (abbreviated as PA14 and PAO1, respectively) impart an advantage during the early stages of biofilm formation (i.e., initial colonization of a clean surface), we used microfluidic devices in which bacteria can be monitored as they attach and grow on glass substrata. To allow for visualization of each strain by widefield or confocal fluorescence microscopy, constitutive fluorescent protein expression constructs were introduced, with a single copy of GFP (ex500/em513) and two copies of *mKO* (ex548/em559) inserted at the *att* site of PAO1 and PA14, respectively (33).

To assess competition during early biofilm formation, PAO1 and PA14 were inoculated into microfluidic chambers in a 1:1 ratio and imaged over 7 hours of flow in a buffered minimal medium containing 1.0 mM K_2_HPO_4_, 0.6 mM MgSO_4_ and 0.4% arginine. We normalized the number of cells for each strain to the number of cells at the start of the assay (0h) and measured the fold change in cell number over time. PAO1 showed a significant advantage in biofilm growth over PA14 as early as four hours after surface inoculation; after 7h of incubation, the number of PAO1 cells was ∼2.5-fold greater than that of PA14 (**Fig. 1A, B, C**).

**Figure 1:**
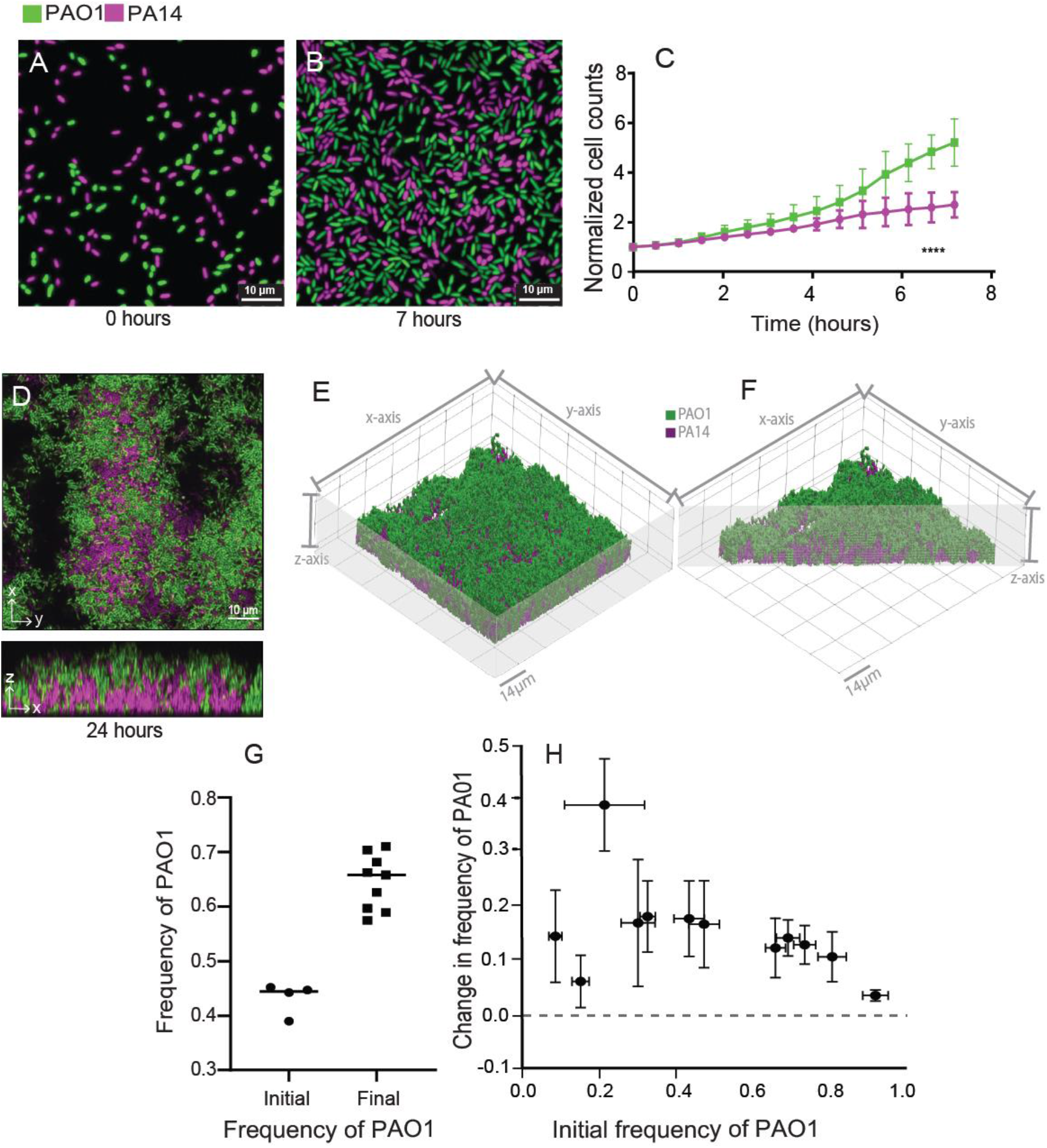
PA14-PAO1 dual strain biofilm dynamics. (**A**,**B**,**D**) Representative widefield fluorescence (A,B) and confocal images (D) of PA14-PAO1 dual strain biofilms at the indicated time. The same color scheme for PAO1 (green) and PA14 (magenta) is used in all panels here, and in all figures in the manuscript. **(C)** Quantification of early biofilm formation (0-7 hrs). N = 3, **** P< 0.0001 after 5.5 hr. (**E**) 3-D rendering of the PA14-PAO1 dual strain biofilm.

If the observed advantage of PAO1 during biofilm competition was due to enhanced growth rate or antagonistic interactions with PA14 via diffusible secreted factors, one would expect a similar competitive outcome in mixed liquid culture conditions. To test this possibility we grew planktonic PAO1 and PA14 in glass tubes containing sterile biofilm medium either in mono-culture or 1:1 co-culture. PAO1 and PA14 grew equally well when cultured separately (**Fig. S1A**), and neither strain outgrew the other in mixed planktonic cultures (**Fig. S1B**). Notably, the CFU counts of the strains in a co-culture were lower by equal measures than their respective CFU counts when grown alone; this indicates neutral competition in which the two strains compete for limited resources in mixed liquid co-culture, but neither has an advantage over the other. These results demonstrate that the advantage in biofilm competition for PAO1 is not due to higher basal growth rate but rather to other root causes.

Biofilms often exhibit complex architectures that can vary significantly in mixed strain or mixed species contexts (34, 35); in light of these observations we next assessed the spatial structure of PAO1 and PA14 in biofilm co-culture. To monitor these dynamics, PAO1 and PA14 were grown together in microfluidic chambers for 24h and then imaged by confocal microscopy. Consistent with the pattern observed in early biofilms inoculated with a 1:1 ratio of PAO1 and PA14 (**Fig. 1A**,**1B**), after 24h of growth PAO1 showed a significant increase in relative abundance compared to PA14 (**Fig. 1D**,**1G**). Z-projections of confocal image stacks revealed that PAO1 grows across the top of the PA14 cell clusters (**Fig. 1D**, bottom panel; **Fig. 1E**,**1F**), an observation consistent with a previous report from Diggle and colleagues (36).

The above data provide evidence that when grown in the biofilm context, PAO1 has an early fitness advantage against PA14 and can ultimately dominate the population. Thus far, a 1:1 inoculation ratio of PAO1 to PA14 was used to assess dual strain population dynamics. This condition does not account for the possibility of frequency-dependent competition, in which the favored strain may depend on the initial ratio of the two. To test if the competitive advantage of PAO1 was frequency-dependent, we varied the starting PAO1 frequency from 0.1 through 0.9 and measured the change in frequency of PAO1 versus PA14 after 24h. Regardless of the starting frequency, PAO1 consistently increased in relative abundance (**Fig. 1H, Fig. S2A**,**S2B**). We also tested whether starting density would affect this dynamic and found that irrespective of the tested starting densities, PAO1 was able to increase in frequency in a dual-strain biofilm (**Fig. S2C**). We conclude from these experiments that under flow in the conditions used here, PAO1 outcompetes PA14 in a frequency- and density-independent fashion.

Rendering is performed as detailed in the Materials and Methods. (**F**) Cut-away 3-D rendering of the same biofilm in panel (E), showing its internal structure. (**G**) Initial (0 h) and final (24 h) frequency of PAO1 in a dual strain PA14-PAO1 biofilm grown at 37°C. (**H**) Change in frequency of strain PAO1 as a function of its initial frequency in dual strain biofilms of wild-type PAO1 and PA14. Initial frequencies (n=4 for each data point) and changes in final frequency (n=9 for each data point) after 24 hours at 37°C are plotted. Error bars represent standard deviation.

### Psl is required for PAO1 to outcompete PA14 in dual strain biofilms

We have provided evidence that PAO1 robustly outcompetes PA14 when both are grown together in biofilms. To clarify the mechanism of these dynamics, we attempted to alter the competition outcome (PAO1 dominance) by reducing the ability of PAO1 to compete or genetically manipulating PA14 to enhance its ability to produce biofilms. *P. aeruginosa* produces three extracellular polysaccharides (EPS) known to facilitate biofilm formation: Pel, Psl and alginate (37). Alginate is broadly conserved in pseudomonads but only conditionally expressed in PAO1 and PA14 during periods of stress (38–40). PAO1 and PA14 both produce Pel, while Psl is unique to PAO1 (41, 42). Previous work has also shown that Psl is a cooperative resource among secreting cells, and cells that do not produce it were excluded and outcompeted in a PAO1 background (36). We hypothesized that Psl provided an advantage for PAO1 not afforded to PA14 and tested a PAOl mutant with a clean deletion of the *psl* promoter (referred to as PAO1 Δ*psl* here) against PA14 in a biofilm. Compared to the WT PAO1 strains, the PAO1 Δ*psl* mutant exhibited a significant decrease in relative abundance in competition with PA14 after 24h (**Fig. 2, Fig. S3A**,**B**).

**Figure 2:**
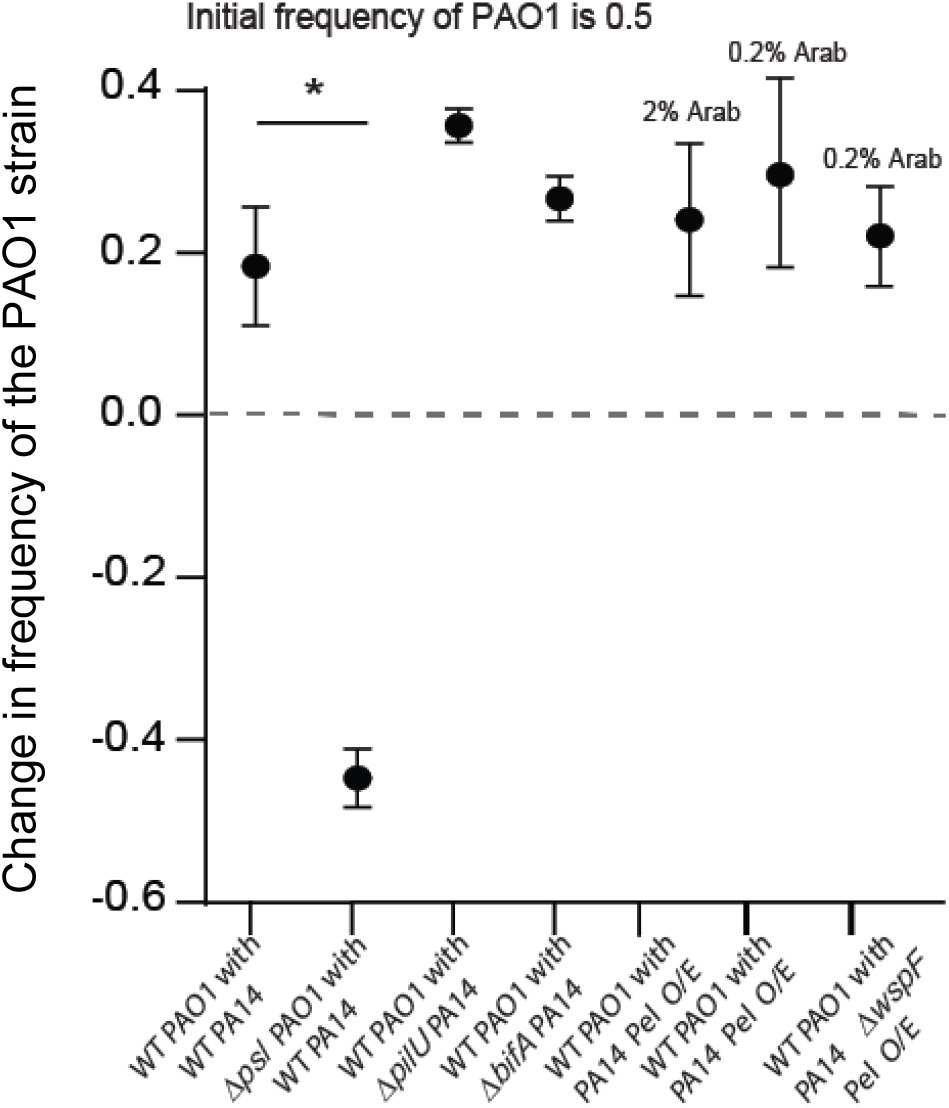
Biofilm mutant analysis. Change in frequency of WT PAO1 and the PAO1 Δ*psl* mutant in the biofilm with WT PA14, as well as change in frequency of WT PAO1 in a dual strain biofilm with PA14Δ*pilU*, PA14 Δ*bifA* and Pel over-producing strains. Initial frequency of all strain combinations was 0.5 (n=5-9). Error bars represent standard deviation; Mann-Whitney tests were used for all pairwise comparisons, * denotes p=0.0010, and all other comparisons were not significant.

We next used a variety of strategies to enhance PA14 biofilm formation and asked whether these altered PA14 strains could better compete versus PAO1. First, we took advantage of the observation that type 4 pili (T4P) have been shown to mediate initial attachment by *P. aeruginosa* (14), and recent work showed that manipulation of T4P functions could enhance surface commitment by PA14 (32). Specifically, we tested if the PA14 Δ*pilU* mutant, a strain that is hyper-piliated, shows high constitutive cAMP signaling and rapidly colonizes a surface, would impart a competitive advantage when grown in a biofilm with PAO1. The PA14 Δ*pilU* mutant was, similarly to wild-type PA14, outcompeted by PAO1 (**Fig. 2)**.

Next, having shown that Psl plays an important role in the ability for PAO1 to compete against PA14, we reasoned that PA14 might gain a competitive advantage against PAO1 through the induction of PA14’s endogenous EPS systems, Pel. To test this hypothesis, we first used a Δ*bifA* mutant, which we showed previously displays increased c-di-GMP signaling and enhanced Pel polysaccharide production (43), but the PA14 Δ*bifA* mutant was not any more successful against PAO1 than WT PA14. As an alternative approach we next used a strain that expresses the *pel* operon under the control of an arabinose-inducible promoter (designated Pel O/E). Interestingly, while induction of *pel* expression with 2% or 0.2% arabinose increased the biovolume of this strain in a monoculture biofilm (**Fig. S3C**), the competitive fitness of the Pel O/E strain induced with high and low arabinose was not enhanced when grown with PAO1 (**Fig. 2**). Finally, we used a variant of the Pel O/E strain which also over produced c-di-GMP due to a mutation in the *wspF* gene (designated Pel O/E Δ*wspF*); again, this strain when induced with 0.2 arabinose did not show enhanced fitness compared to the PA14 parent strain (**Fig. 2**) despite enhanced biomass in monoculture (**Fig. S3C**). Taken together, these data show that Psl production is a key factor allowing PAO1 to outcompete PA14, and that enhancing PA14 biofilm production via several different strategies is unable to overcome the advantage displayed for PAO1 during biofilm formation.

### PAO1 dominates in an environment marked by perturbation

It is possible that although PA14 is outcompeted by PAO1 within a growing biofilm, PA14 might be better suited to dispersal to new locations for future biofilm growth. Therefore, we next asked if PAO1 still dominated in an environment marked by perturbation. To test this idea, we grew a 1:1 mixture of PA14 and PAO1 in microfluidic chambers under flow for a fixed time (either 20h or 3h) after which we introduced a disturbance event. For each such event, the outflow tube from the first microfluidic chamber was attached to a second previously uncolonized, clean chamber and the biofilm effluent was used to seed this microfluidic chamber for 2 hrs. The goal was to simulate the natural transition of *P. aeruginosa* from an existing biofilm to a new environment with an intervening planktonic phase (**Figure 3A**, top).

**Figure 3:**
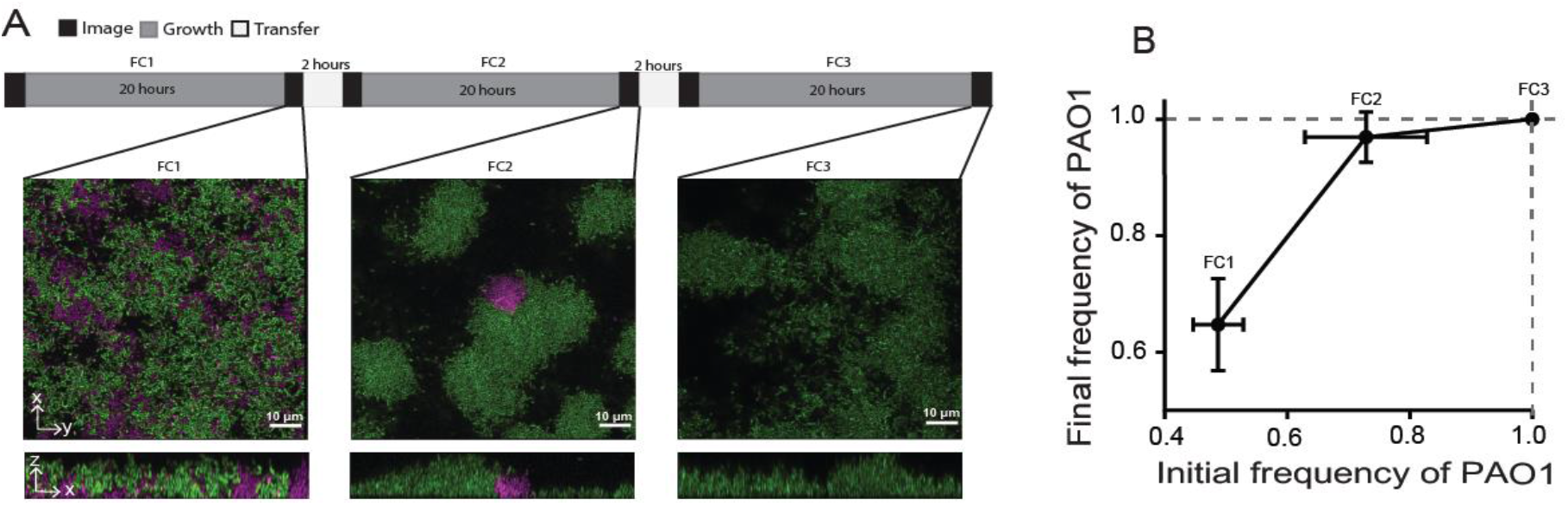
PAO1 dominates during a perturbation regime. (**A**) Graphical summary of dispersal experiment regime (top) and representative images (bottom). (**B**) The frequency dynamics of PAO1 in dispersal experiments through different chambers (n = 3). All error bars shown are standard deviation.

In dual-strain biofilms that were inoculated 1:1 and grown undisturbed (also shown above), PAO1 accounted for ∼75% of the population at the end of the 20-h incubation (**Fig. 2**,**3A**,**3B**). In comparison, with disturbance events at 20 h intervals, the percentage of PAO1 drastically increased compared to PA14 in chamber 2 (after 1 disturbance) to about 97% of biofilm biovolume. After the second disturbance, PAO1 comprised ∼100% of the biofilm by chamber 3 (**Fig. 3A, 3B**). These data indicate that the disturbance regime still favors PAO1 versus PA14 when the two strains compete to form biofilms in successive environmental patches following dispersal.

To test if this observed equilibrium in chamber 3 (after two disturbances) was a function of PAO1 having overgrown PA14 (**Fig. 1D**,**1E**,**1F**), we repeated this disturbance assay in 3-h biofilms; at this early time point PAO1 has not yet had sufficient time to overgrow PA14. As there were relatively few cells attaching at 3 hrs, quantifying population frequencies was not possible post dispersal as was done for the 20 hr perturbation experiments, so we instead calculated the frequencies observed in chambers two and three after 3h of growth (Fig 3B), and then then inoculated new chambers with the same frequency but at a ∼500-fold higher density. Similar to the trend in 20-h disturbance experiments we saw an increase in PAO1 frequency in the subsequent chamber at ∼60% in chamber 2 and ∼70% in chamber 3 (**Fig. S4**), again consistent with PAO1 outcompeting PA14 in a fluctuating environment requiring re-colonization of new surfaces.

### PA14 is more proficient at invading pre-formed biofilm of PAO1

The data above suggest that PAO1 can dominate PA14 when in competition to colonize a new surface, or under conditions of repeated environmental perturbation. These observations left open the question as to what condition or conditions might PA14 dominate over PAO1, and we next hypothesized that while PAO1 excels in colonizing new environments and competing in them thereafter, perhaps PA14 has an advantage when colonizing areas in which pre-existing biofilms reside.

To test this hypothesis, we grew a biofilm of one of the strains, which we refer to as the “ resident strain”, in a microfluidic chamber. After 12h of growth of the resident strain, we introduced the second strain, referred to as “ invader”, for 4 hours to assess its ability to colonize and integrate into the resident biofilm.

By visual inspection alone it was evident that PAO1 showed minimal invasion into resident PA14 biofilms (**Fig. 4A)**, while PA14 was considerably more proficient at invading pre-formed biofilms of PAO1 (**Fig. 4B**). To quantify invasion efficiency by PAO1 and PA14, we normalized the fluorescent intensity for each strain at the start of widefield fluorescence imaging (0h) and measured the fold change in signal intensity over 4h of the invasion assay. PA14 invades resident PAO1 biofilms rapidly after introduction to the chamber (**Fig. 4B**,**4C**), and by the end of the assay, PA14 invading biovolume was ∼3 orders of magnitude higher than that of PAO1 invading a resident PA14 biofilm, as measured by confocal microscopy (**Fig. 4D)**.

**Figure 4:**
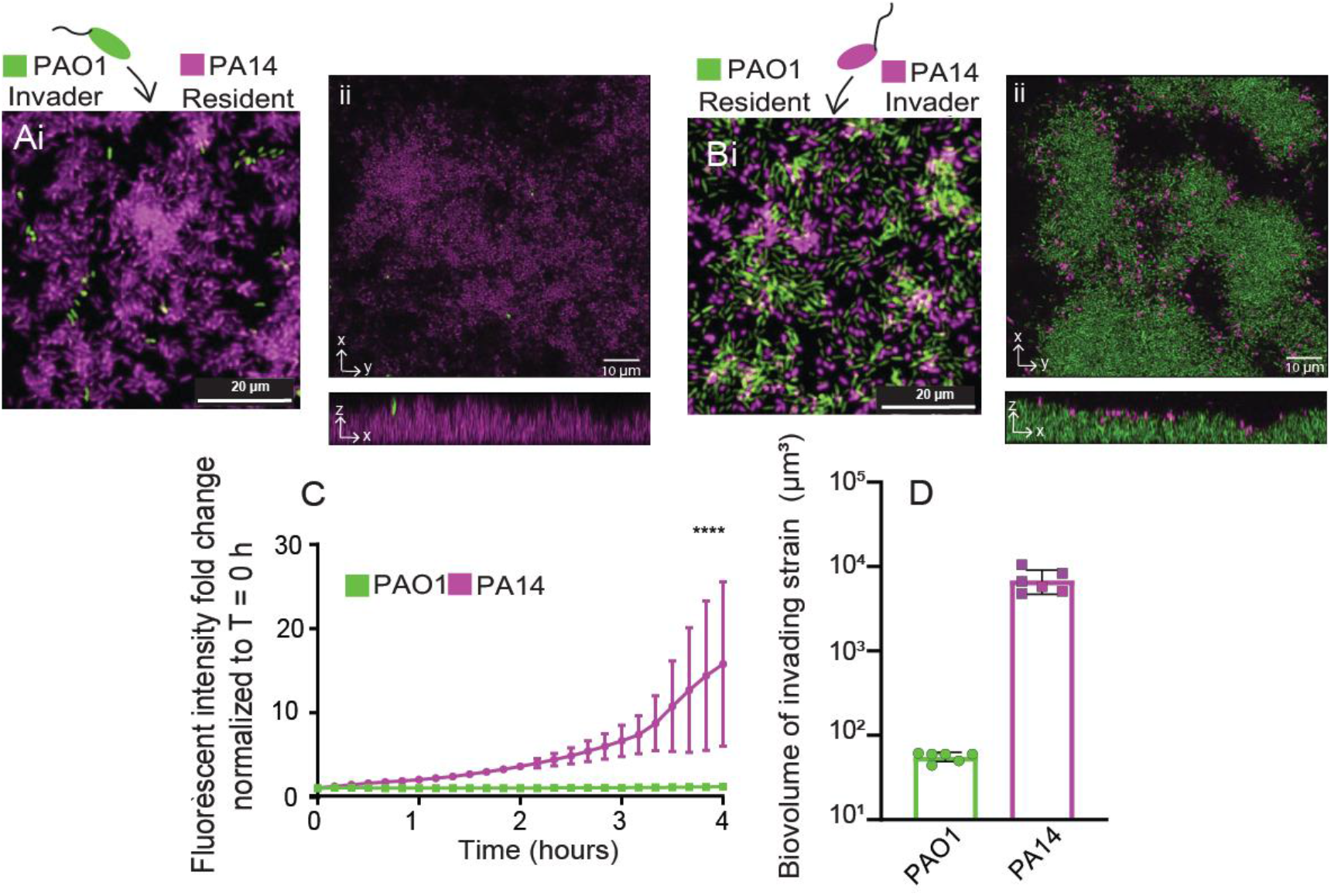
Invasion dynamics of resident biofilms. (**A**) Resident biofilms of PA14 were grown for 12h and invaded by PAO1 for 4h (i) widefield fluorescence image (ii) confocal images with x-y optical section above and z projection below. (**B**) Resident PAO1 biofilms were invaded by PA14. (i) widefield fluorescence image (ii) confocal images with x-y optical section above and z projection below. (**C**) The invasion efficiency of PA14 (purple) and PAO1 (green) was measured over time by normalizing the change in signal intensity from the start of the assay through 4h of invasion. n = 3, **** P< 0.0001 after 3h by ANOVA with a Sidak post-test. (**D**) The total invading strain biovolume of PAO1 and PA14 at the end of the invasion assay (n = 6).

### PA14 maintains surface coverage relative to PAO1 after starvation

The studies above assess the ability of PAO1 and PA14 to compete in the context of colonization. Given that the supply of nutrients is transient and irregular in most natural environments, we decided to investigate how PAO1 and PA14 would react to nutrient depletion. To address this question, we grew dual strain biofilms for 12 hour and then changed the influent flow to a carbon-free biofilm medium for 4 h. We observed that strain PA14 had higher persistence with only a ∼5% reduction in surface coverage of cells when carbon-free medium was introduced (**Fig. 5A, 5B**). By contrast, PAO1 exhibited an ∼60% decrease in surface area coverage over the same time period after influent exchange to carbon-free medium (**Fig. 5A, 5C)**, suggesting that PAO1 is more likely to disperse under nutrient-limited conditions.

**Figure 5:**
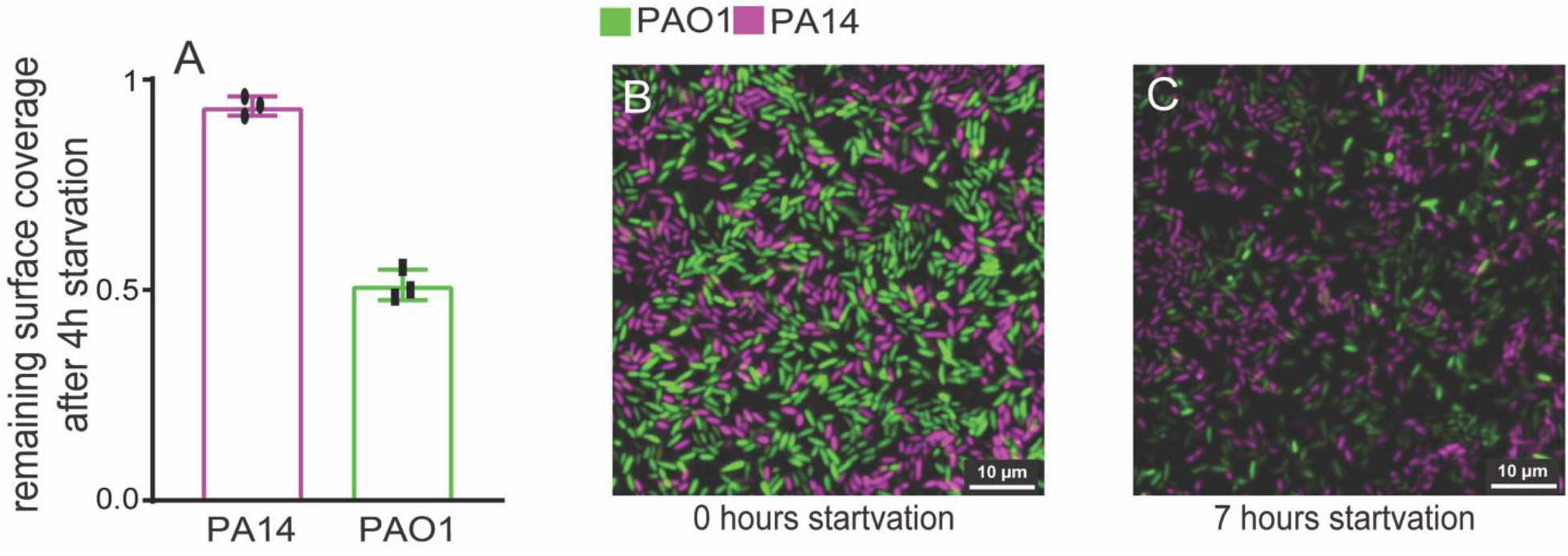
Starvation tolerance of PAO1 and PA14 biofilms. Co-culture biofilms of PAO1 and PA14 were grown for 12h and deprived of a carbon source (arginine) for 4h. (**A**) The surface area occupied by PAO1 and PA14 after 4h starvation was measured as a fraction to the surface area prior to starvation. Error bars represent standard deviation, n=3. Representative widefield fluorescence microscopy images of co-culture PAO1 and PA14 biofilms prior to starvation (**B**) and after 4h starvation (**C**).

## Discussion

Here we investigated reference strains *P. aeruginosa* PAO1 and PA14 for their ability to compete with each other in the context of biofilm growth. Prior work has provided evidence that PAO1 uses the Wsp system to quickly commit to surfaces through a processive mode of attachment, while PA14 uses the Pil-Chp signaling system to mediate non-processive surface engagement (19). We provide evidence that in microfluidic biofilm culture conditions, PAO1 quickly outcompetes and overgrows PA14 in a density and frequency-independent manner. PAO1 and PA14 were also found to spatially segregate on 10 micron length scales, with PA14 limited to the substratum and PAO1 over-growing PA14. The competitive advantage of PAO1 is specific to the biofilm mode of growth, as PAO1 did not outgrow PA14 in planktonic mono- or co-culture. Finally, using a regime that mimics environmental perturbation, we show that PAO1 remains competitively dominant and indeed reaches fixation in the population after only two recolonization events.

To understand how PAO1 outcompetes PA14 we tested a Psl mutant of PAO1 as well as PA14 mutants with increased biofilm formation or elevated levels of Pel and found that only through loss of Psl in the PAO1 background did PA14 outcompete PAO1 in biofilm co-culture. Interestingly, while induction of Pel in PA14 led to increased biofilm biomass accumulation in monoculture, increased Pel production did not translate into better competitive ability in mixed culture, a result which suggests that Psl may provide a unique advantage for PAO1 in biofilm environments. The importance of Psl has also been observed in polymicrobial biofilms; consistent with our data, the loss of Psl resulted in a significant decrease in the biovolume of PAO1 when grown with other species such as *Pseudomonas protegens* and *Klebsiella pneumoniae* (44).

We did note two conditions where PA14 seemed to be superior in competition against PAO1 in biofilm co-culture, namely in colonizing a preformed biofilm of PAO1 and in retaining biomass in place in the face of nutrient deprivation. PA14 was better able to invade pre-formed biofilms of PAO1 by ∼3 orders of magnitude than vice versa, suggesting that PA14 may be better suited to exploiting previously colonized environments, while PAO1 is superior in competition for space and resources on unoccupied surfaces. Furthermore, PA14 may be more suited to retaining a grip on space it has occupied when nutrients run low, while PAO1 is more inclined to disperse. The relative of advantage of staying in place under starvation conditions depends on the prevailing environmental conditions: if local nutrient supply fluctuates, then remaining in place may be the better strategy, but if nutrient supply does not return once depleted, then dispersal will be optimal. Recent genomic analysis has found that strains similar to PA14 predominant among CF-derived isolates, while the PAO1-like strains are more likely to be encountered among environmental isolates (45). We speculate on the basis of our results that PA14 has evolved a surface occupation strategy best suited to taking advantage of previously colonized surfaces and commitment to staying in place under fluctuating nutrient conditions, while PAO1 optimizes rapid exploitation of unoccupied surfaces followed by rapid dispersal under nutrient limitation to find new locations for future growth. Thus, as we had predicted in our previous study (19), the differential colonization strategies of PAO1 and PA14 do indeed provide competitive advantages in different contexts.

## Materials and Methods

### Microfluidic device assembly

The microfluidic devices were made by bonding polydimethylsiloxane (PDMS) chamber molds to size #1.5 cover glass slips (60mm × 36mm [LxW], Thermo-Fisher, Waltham MA) using standard soft lithography techniques (46). Each PDMS mold contained 4 chambers, each of which measured 3000 µm⨯500 µm ⨯ 75 µm (LxWxD). To establish flow in these chambers, medium was loaded into 1 mL BD plastic syringes with 25-gauge needles. These syringes were joined to #30 Cole-Parmer PTFE tubing (inner diameter 0.3 mm), which was connected to pre-bored holes in the microfluidic device. Tubing was also placed on the opposite end of the chamber to direct the effluent to a waste container. Syringes were mounted to syringe pumps (Pico Plus Elite, Harvard Apparatus), and flow was maintained at 0.1 µL per min for all experiments.

### Biofilm growth

Overnight cultures of *P. aeruginosa* strains were grown at 37°C shaking in lysogeny broth (LB) prior to the start of biofilm experiments. Cultures were normalized to an OD_600_ of 1 in KA biofilm (47) medium containing 50mM Tris-HCl (pH 7.4), 0.6 mM MgSO_4_, 1.0 K_2_HPO_4_ and 0.4% arginine. If dual strain biofilms were assessed, equal volumes of cultures adjusted to OD_600_ of 1 were mixed and used as the inoculum for the microfluidic chamber (completely filling its inner volume), and then the bacteria were allowed to rest for 1 h at room temperature to permit cells to attach to the glass surface. For the experiments with varied initial frequencies, after cultures of each strain were adjusted to an OD_600_ of 1, ratios of the two cultures were added to obtain the desired frequency prior to inoculation. After resting for 1 hour to allow bacterial attachment, the devices were run at 0.5 µL per min at 37°C and imaged by widefield or confocal microscopy (see below) at time intervals that varied per experiment as noted in the main text. At every sampling time point, images were acquired from nonoverlapping locations within each biofilm chamber. All experiments were repeated with at least 3 biological replicates with 3 or more technical replicates on different days. Total replicates for each experiment are noted in the figure legends for each data set in the text and Supplementary Information.

For the perturbation studies, a dual strain biofilm (1:1 initial frequency of PA14: PAO1) was used to seed a biofilm for either 24 hours or 3 hours, and a 0.5-cm length of tubing was connected to the outlet channel. At every sampling time point, images were acquired from nonoverlapping locations within each biofilm chamber. The outlet chamber was then allowed to seed a new chamber; this process was repeated two times for the 24 hour incubation studies. For the 3-hour protocol, because of low cell numbers attaching and the inability to quantify those low biovolumes, we conducted the disturbance protocol and tabulated the frequency of the two strains in the newly colonized chamber. We then inoculated a new chamber with the same frequency with about 500-fold increased density to allow sufficient biomass to accumulate to determine the relative frequency of each strain.

For invasion experiments, we grew the resident biofilm for 12 h at a media flow rate of 0.5 µL per min at 37°C. After which we introduced the invading strain (adjusted to an OD_600_ = 1) at the same flow rate for 4 hours. At every sampling time point, images were acquired from nonoverlapping locations within each biofilm chamber.

### Microscopy and image analysis

Biofilms in the microfluidic chambers were imaged using a Zeiss LSM 880 confocal microscope with a 40x/1.2NA or 10x/0.4NA water objective. A 543-nm laser line was used to excite mKO-κ, and a 594-nm laser line was used to excite mKate2. A 458-nm laser line was used to excite Wisteria floribunda lectin stain in the case of Pel quantification experiments. All quantitative analysis of microscopy data was performed using BiofilmQ (48). 3-D renderings of biofilms in Figure 1 and Figure 4 were generated using Paraview.

### Statistics

All statistical analyses were performed in GraphPad Prism. All reported pairwise comparisons were performed using Wilcoxon signed-ranks tests, and multiple comparisons were performed by Wilcoxon signed-rank tests with Bonferroni correction. All error bars indicated standard deviation unless otherwise noted.

## Acknowledgements

This work was supported by funding from the NIH (R37-AI83256-06) to GAO. CDN was supported by the Cystic Fibrosis Foundation (STANTO19RO), NSF (MCB-1817342) and NIH (P30-DK117469).

